# Lineage-specific oncogenes drive growth of major forms of human cancer using common downstream mechanisms

**DOI:** 10.1101/2022.09.27.509636

**Authors:** Otto Kauko, Mikko Turunen, Päivi Pihlajamaa, Antti Häkkinen, Rayner M. L. Queiroz, Mirva Pääkkönen, Sami Ventelä, Massimiliano Gaetani, Susanna Lundström, Antonio Murgia, Biswajyoti Sahu, Johannes Routila, Heikki Irjala, Julian L. Griffin, Kathryn S. Lilley, Teemu Kivioja, Sampsa Hautaniemi, Jussi Taipale

**Affiliations:** Applied Tumor Genomics Program, University of Helsinki, Biomedicum, P.O. Box 63 (Haartmaninkatu 8), FIN-00014 University of Helsinki, Finland; Department of Biochemistry, University of Cambridge, U.K.; Turku Bioscience, University of Turku and Åbo Akademi University, Finland; Systems Oncology Program, University of Helsinki, Biomedicum, P.O. Box 63 (Haartmaninkatu 8), FIN-00014 University of Helsinki, Finland; Cambridge Centre for Proteomics, University of Cambridge, U.K.; Department for Otorhinolaryngology - Head and Neck Surgery, University of Turku and Turku University Hospital, Finland; Department of Medical Biochemistry and Biophysics, Karolinska Institutet, Sweden; Centre for Molecular Medicine Norway, University of Oslo, Oslo, Norway; Rowett Institute, University of Aberdeen, U.K.; Department of Computer Science, P.O. Box 68, FIN-00014 University of Helsinki, Finland

## Abstract

Mutations in hundreds of genes have been associated with formation of human cancer, with different oncogenic lesions prevalent in different cancer types. Yet, the malignant phenotype is simple, characterized by unrestricted growth of cells that invade neighboring healthy tissue and in many cases metastasize to distant organs. One possible hypothesis explaining this dichotomy is that the cancer genes regulate a common set of target genes, which then function as master regulators of essential cancer phenotypes, such as growth, invasion and metastasis. To identify mechanisms that drive the most fundamental feature shared by all tumors – unrestricted cell proliferation – we used a multiomic approach to identify common transcriptional and posttranslational targets of major oncogenic pathways active in different cancer types, and combined this analysis with known regulators of the cell cycle. We identified translation and ribosome biogenesis as common targets of both transcriptional and posttranslational oncogenic pathways. By combining proteomic analysis of clinical samples with functional studies of cell cultures, we also establish NOLC1 as a key node whose convergent regulation both at transcriptional and posttranslational level is critical for tumor cell proliferation. Our results indicate that lineage-specific oncogenic pathways commonly regulate the same set of targets important for growth control, revealing novel key downstream nodes that could be targeted for cancer therapy or chemoprevention.

## INTRODUCTION

Mutations in more than 500 genes have been causally linked to cancer^1, 2^. Several genes exist that are commonly mutated in many different cancer types, such as p53, TERT, ATM, and CDKN2A^3, 4^. These genes are implicated in processes such as maintaining DNA integrity and progression through cell cycle checkpoints. However, most known cancer genes exhibit some degree of tissue specificity. Notably, genes encoding proteins involved in growth signal transduction, including members of the Wnt, Hedgehog and tyrosine kinase/RAS/PI3K signaling pathways exhibit some of the highest observed mutation frequencies in select cancers while not being mutated at observable rates in others^5^.

Growth signal transduction promotes the expression of genes needed for cell proliferation. Some signal transduction pathways, such as Wnt and Hedgehog activate specific transcription factors with well-known target genes. Activation of tyrosine kinase/RAS/PI3K signaling can also regulate activity of specific transcription factors, but the majority of their downstream targets have an unknown function^6^. It is thus unclear whether transcriptional regulation is the main shared outcome of phosphorylation signaling, or is its growth-promoting effect also transduced more directly via effects on protein activity levels, e.g. via upregulation of metabolism and translation^7^.

Established genetic methods have inherent limitations in their capacity to systematically identify all targets for mechanism-based cancer therapy and prevention. Somatic cancer genetics identifies genes that cause cancer, not proteins whose inactivation specifically kills cancer cells. In addition, while a somatic mutagenesis “screen” may be saturating with respect to individual genes and amino-acids, it lacks the power to detect combinations of more than ∼ 10 mutations. This leads to failure to detect large protein complexes and pathways, wherein increase in levels of all proteins are needed for increase in activity. In such a case, each member of a pathway or protein complex can still be individually necessary for the oncogenic activity, and thus represent a potential druggable target.

Another highly successful genetic approach, genome-wide association study (GWAS), identifies variants that increase or decrease cancer risk^8^. This method has high sensitivity but can access only a very small subset of all theoretically possible variants. GWAS primarily detects variants that affect protein expression levels; this leads to a bias against proteins whose activity is controlled at other levels such as posttranslational modification, degradation, or allostery. For example, many genes that are critically important for cancer, such as Ras and p53 do not show up prominently in GWAS. Given that genetic approaches based on both somatic mutation and inherited risk have innate limitations in detecting potential drug targets, we hypothesize that a variety of presently unknown mechanisms exist whose activity could be targeted to treat or prevent cancer (**Fig. 1**).

**Figure 1.**
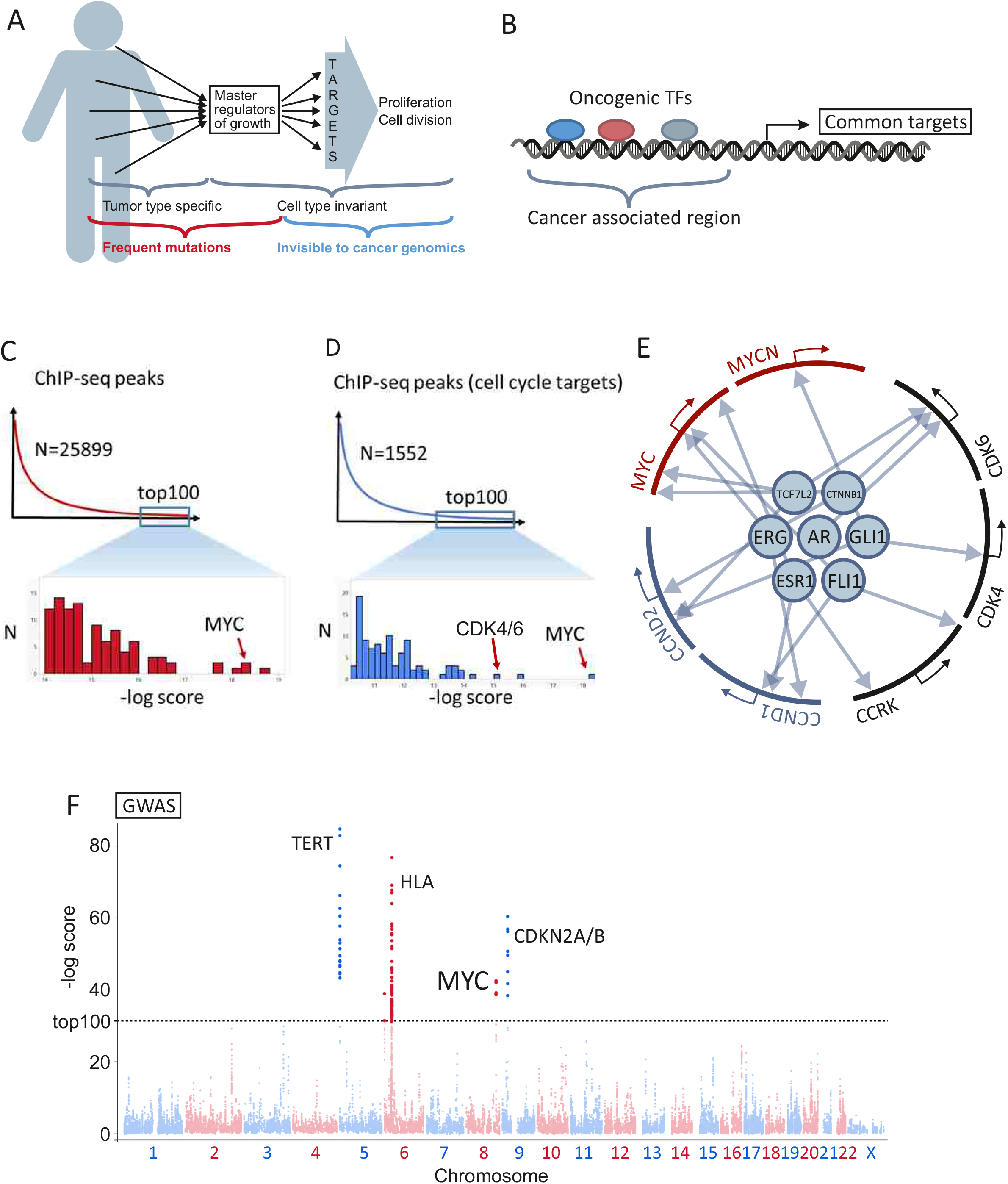
Transcriptional regulation of cell growth. **A)** Proposed model based on the hypothesis that multiple signaling pathways and oncogenes drive the cell cycle by targeting a limited set of downstream genes which function as master-regulators of growth. **B)** Oncogenic transcription factors (TFs) exhibit convergent functions by binding to the same enhancer elements to engage shared downstream genes. **C)** Shared target genes of oncogenic TFs were identified by chromatin immunoprecipitation followed by sequencing (ChIP-seq). Top-scoring common target genes of oncogenic transcription factors based on the number and heights of different ChIP-seq peaks and their proximity to the transcription start site. Scoring-scheme for panels C,D, and F is described in detail in the Supplementary Methods. **D)** Top-scoring common target genes are shown with the restriction that they are involved in cell cycle regulation in human cells^19^ or that their orthologs affect the cell cycle in *Drosophila*^*20*^ **E)** Results of the ChIP-seq analysis shown for the enhancers of selected paralog groups. TF Binding site positions are indicated with an arrow with respect to transcription start site (TSS). Red: MYC genes, Black: CDK4/6, CCRK. Blue: D-type cyclins **F)** Scoring of the cancer associated SNPs in GWAS Catalogue. Top 100 highest scoring SNPs are indicated with darker color above the dashed line. Highest scoring SNPs are from four loci corresponding to TERT, HLA antigen, MYC, and CDKN2A/B.

We propose here a model that would explain both the complexity of the cancer genotype and the tissue-type specificity of oncogenes. In this model, different signaling pathways that drive major forms of human cancer in different tissues converge on the same functional output – unrestricted cell proliferation. This occurs because the gene regulatory network that controls growth has an hourglass shape. The large number of tissue-specific oncogenic proteins activate few common master regulators, which then control a large number of proteins that contribute to the downstream transcriptional and posttranslational processes necessary for upregulation of metabolism and cell proliferation (**Fig. 1A**).

## RESULTS

### Oncogenic transcription factors from major forms of human cancer regulate MYC

As targeting oncogenic drivers in any particular tumor results in a large number of changes in gene expression and posttranslational modifications, we reasoned that generation of a large number of datasets, and focusing on the common features across multiple of them would enable us to focus on a more limited set of downstream mechanisms that are common to multiple major forms of human cancer. To ensure that all data was comparable, we generated all the primary input datasets to the analysis in house.

We first set out to identify common transcriptional targets of oncogenic TFs. We reasoned that convergence of oncogenic transcriptional mechanisms would most likely occur via enhancer elements (**Fig. 1B**). Key master regulators would contain multiple enhancers, and the tissue-specificity of oncogenes would be explained by the fact that a given oncogenic TF activates a particular enhancer by collaborating with tissue-specific factors^9^.

To identify direct targets of oncogenic TFs, we used ChIP-seq to detect binding of estrogen receptor in breast cancer ^10^, androgen receptor and ERG in prostate cancer^11^, Tcf4 and β-catenin in colorectal cancer ^12^, GLI1 and PAX3 in rhabdomyosarcoma ^13-15^, and FLI1 in Ewing’s sarcoma^16^ (**Table S1**). The transcription factors analyzed have all been previously shown to be critical for the formation of the tumors, and involved in a substantial fraction of all cases of the respective forms of cancer (see Refs. ^10-18^).

The number of peaks identified for each factor is indicated in **Fig. S1A**. We also performed expression profiling analyses to identify genes whose expression is affected by the transcription factors (**Fig. S1B**). These results were incorporated into a gene-regulatory network in Cytoscape, which is modeled as follows: There are four classes of nodes: transcription factors, ChIP-seq peaks, cancer associated regions (from GWAS) and target genes. There is an edge from a TF node to all its peak nodes, and there is an edge from a peak or association node to a target gene node if they are within 500 kb of the target gene. A direct edge between a TF and a target gene is drawn if a target gene expression is altered after RNAi of the TF. The peak heights and enhancer scores, as well as the distances of the elements from the TSS of the gene are given as attributes to the respective nodes and edges (see **Supplementary Methods** for details). To identify common targets of oncogenic TFs, we implemented a subgraph isomorphism algorithm that searches for paths converging on a single gene as a Cytoscape plug-in.

Using these tools, we drew query networks, and executed the plug-ins to identify all subgraphs from the larger network that were isomorphic to the query. To facilitate visual analysis, a post-processing step was used to merge all subgraphs that contain target genes in the same genomic regions (**Fig. S2**). The target genes were then ranked based on the number of different ChIP-seq peaks and their proximity to the transcription start site (measured as ranked distance; see **Supplementary Methods** for details). Due to gene duplication events, humans often have many genes whose proteins have very similar if not identical functions. Such proteins often are regulated differentially, and could thus be targeted by different oncogenic transcription factors in different tumor types. To address this, we merged paralogous target genes to groups which are likely to have similar activities (see **Supplementary Methods** for details).

Significance of results was assessed by permutation analysis. Initial analysis revealed that the paralog group corresponding to the MYC oncogenes – known targets of many oncogenic signals – was ranked third from all of the paralog groups (**Fig. 1C**). Restricting the set of genes analyzed to known regulators of the cell cycle in human cells ^19^ or that their orthologs affect the cell cycle in *Drosophila* ^*20*^ resulted in identification of MYC and the cyclin dependent kinase CDK4/6/CCRK genes as common targets of lineage-specific oncogenic transcription factors (**Fig. 1D**). Importantly, analysis of the ChIP-seq peaks of the individual oncogenic TFs in the different tumor types showed that the signal was not derived from a single tumor type or an individual TF, but represented broad-based regulation of the master regulatory genes by the oncogenic TFs (**Fig. 1E**).

Similar analysis using only genome-wide association data from diverse cancer types also identified five paralog groups, including MYC genes and components of the CDK4/6 system (CDK inhibitors CDKN2B-D and D-type cyclins) (**Fig. 1F**). In addition, GWAS identified HLA and TERT loci, which were not detected in the ChIP-seq based analysis. This could be because TERT and HLA can affect tumorigenesis indirectly, by causing chromosomal instability and predisposing individuals for infection by tumor-causing viruses, such as human papillomavirus and Epstein-Barr virus. In summary, using two sets of completely orthogonal data, we show that upregulation of the CDK4/6 system and the MYC family of oncogenes are the predominant cell-autonomous outcomes of the activation of lineage-specific oncogenic transcription factors across major forms of human cancer.

### MYC also mediates the major shared transcriptional output of phosphorylation signaling

To identify common transcriptional mechanisms activated by phosphorylation signalling (**Fig. 2A**), we developed a set of cell lines that were either sensitive or resistant to common cytostatic kinase inhibitors. The ten different parental cell lines used represented lung, colorectal and breast carcinomas and chronic myeloid leukemia (CML), and were driven by hyperactivation of phosphorylation signals by mutations in BRAF, RAS, EGFR, ERBB2, PIK3CA and BCR/ABL (**Fig. 2B**). In developing the cell lines, we made use of the redundancy of the phosphorylation signalling, activating known alternate pathways that commonly rescue growth in the presence of a drug targeting a particular kinase^21^.

**Figure 2.**
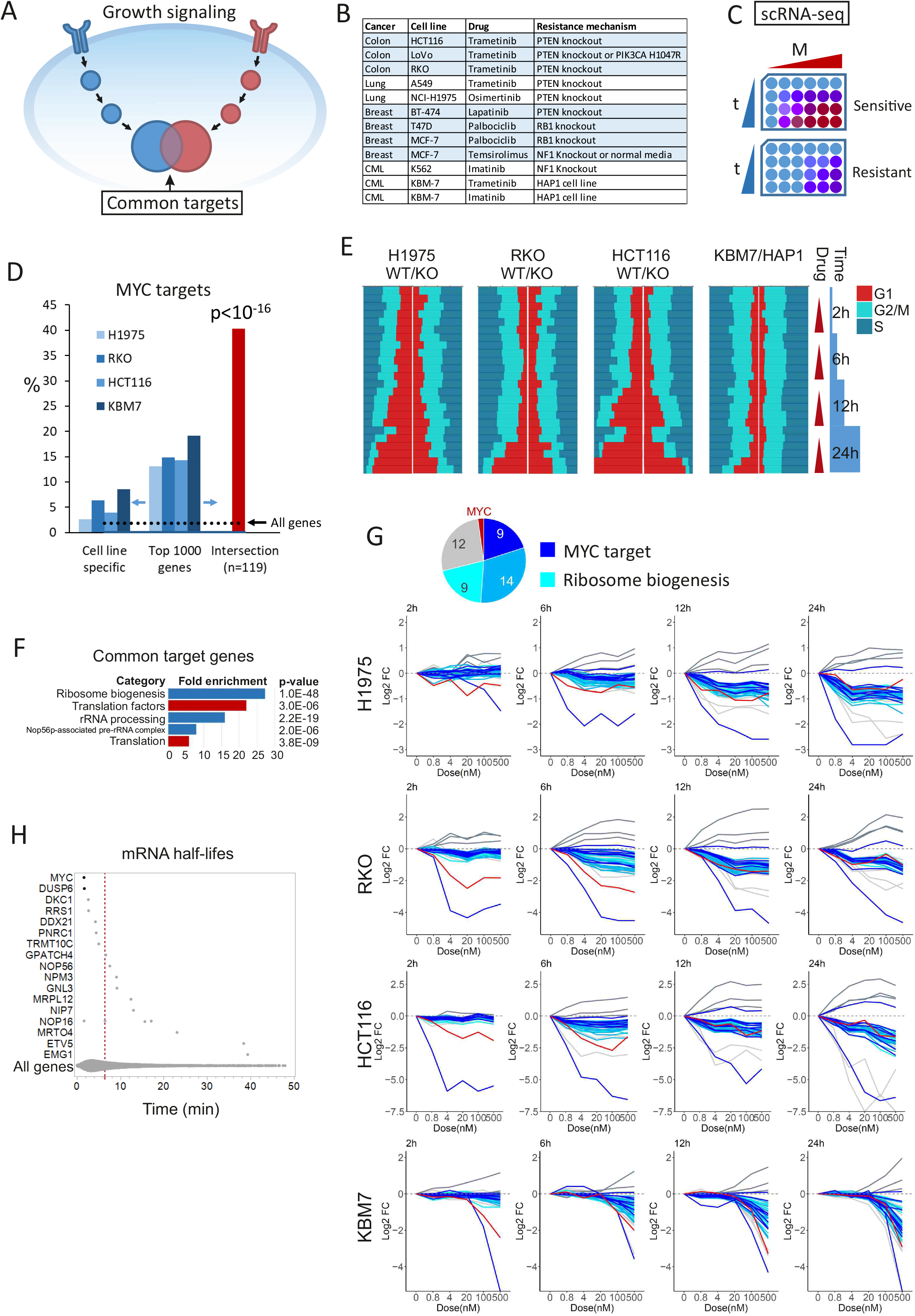
Common transcriptional mechanisms activated by phosphorylation signalling. **A)** Proposed model of phosphorylation signaling pathway convergence based on the hypothesis that the downstream kinases of growth-regulatory pathways phosphorylate (functionally) overlapping sites. **B)** Cell lines, cytostatic kinase inhibitors, and resistance mechanisms in the cell lines generated for this study **C)** Experimental setup for sc-RNAseq. Sensitive and resistant versions of the cell lines were subjected to varying concentrations of the drugs (0-500nM) and sc-RNAseq analysis was performed at 2, 6, 12, and 24 hour time points. **D)** Genes were ranked for each cell line pair based on the explanatory power of drug-induced changes in the sensitive version of the cell line in a multiple regression model. Top genes exhibited significant overlap between different cell lines and were highly enriched in MYC target genes (see Supplementary Methods for details). Fraction of MYC target genes is shown on the Y-axis. In panels D-G, the data were generated from NCI-H1975, RKO, HCT116, and KBM7/HAP1 cells as presented in panel B. **E)** Cell cycle distributions of the cell line pairs. Red: G1, Dark blue: S, Cyan: G2/M. G1 arrest occurs predominantly in the sensitive version of the cell line and at lower drug concentrations. **F)** Enrichment of selected biological categories in the common top ranking genes similar to panel D. Enrichments were calculated with G:profiler^51^ using Benjamini-Hochberg adjustment for multiple hypothesis correction. Database entries for the enriched categories are: Ribosome biogenesis: GO:0042254, Translation factors: WP:WP107, rRNA processing: REAC:R-HSA-72312, Nop56p-associated pre-rRNA complex: CORUM:3055 Translation: GO:0006412 **G)** Expression changes in the common target genes as a function of drug concentration at each time point. Intersection of the Top ranked 500 genes for each cell line is shown for clarity. Red=MYC, Dark blue= MYC target genes, Cyan = Ribosome biogenesis (Gene ontology term 0042254) **H)** mRNA half-lives of the common target genes in panel G. Half-lives were extracted from Chen et al^22^. Median of all genes is shown with a red dashed line

We first treated a set of sensitive and resistant cell lines representing lung and colorectal cancer and CML with cytostatic kinase inhibitors targeting EGFR, MEK, and BCR-ABL. Cells treated with different drug concentrations for different times were labelled with DNA tags (**Fig 2C**; see **Methods**), pooled and subjected to single-cell RNA sequencing (scRNA-seq). Gene expression changes were modeled as a function of drug concentration, treatment duration, presence of resistance mutations, and cell cycle phase. The cell cycle phase was included in the regression model to separate the direct effects of the drugs from the indirect effects caused by arrest of the cell cycle. In the absence of resistance mutations, the cytostatic drugs regulated a highly overlapping set of genes that was heavily enriched in MYC targets (**Fig. 2D**; see **Supplementary Methods**). Drug treatment resulted in G1 cell cycle arrest starting at 12 to 24 h. In the resistant cell lines, cell cycle arrest was incomplete and/or required a higher drug dose **(Fig. 2E**).

Comparison across the cell lines resulted in identification of genes that were regulated prior to the cell cycle arrest in a highly convergent manner in the different cell lines treated with distinct cytostatic drugs (**Table S2**). The genes were heavily enriched in regulators of translation and ribosome biogenesis (**Fig. 2F**). The regulation of MYC preceded the downregulation of its targets, suggesting that most shared transcriptional responses observed after the drug treatments were dependent on MYC (**Fig. 2G**). Ribosome biogenesis factors were regulated with slightly faster kinetics on average than the rest of the MYC targets (visible at the 6h time-point).

Only few of the genes appeared to be regulated independent of MYC. These included DUSP6, ETV4, and ETV5, which were downregulated, and the negative regulator of ribosome biogenesis PNRC1 and two non-coding RNAs (MALAT1 and NEAT1), which were upregulated. The early downregulation of DUSP6 and MYC is most likely due to the corresponding mRNAs having a shorter mRNA half-life than the mRNAs of the other downregulated genes ^22^(**Fig. 2H**). In summary, these results indicate that most transcriptional consequences of cytostatic drugs that are common to multiple cancer types are mediated by downregulation of the MYC oncogene.

### Common phosphorylation targets across human cancer types

In addition to transcriptional output, it is well established that posttranslational regulation has also more direct effects on protein activities. In order to study such non-transcriptional effects of growth signaling, we performed proteomics analyses with ≤2h time points, before the majority of the transcriptional changes or cell cycle effects would take place (**Fig. 3A**). Based on the sc-RNAseq, cell cycle, and cell viability analyses, we selected drug concentrations that only cause cell cycle arrest in the drug-resistant derivative of each cell line. We performed a phosphoproteomics characterization of the cell lines listed in **Fig. 2B**, identifying more than 48,000 phosphopeptides in total (median 19771 per cell line) at <1% FDR.

**Figure 3.**
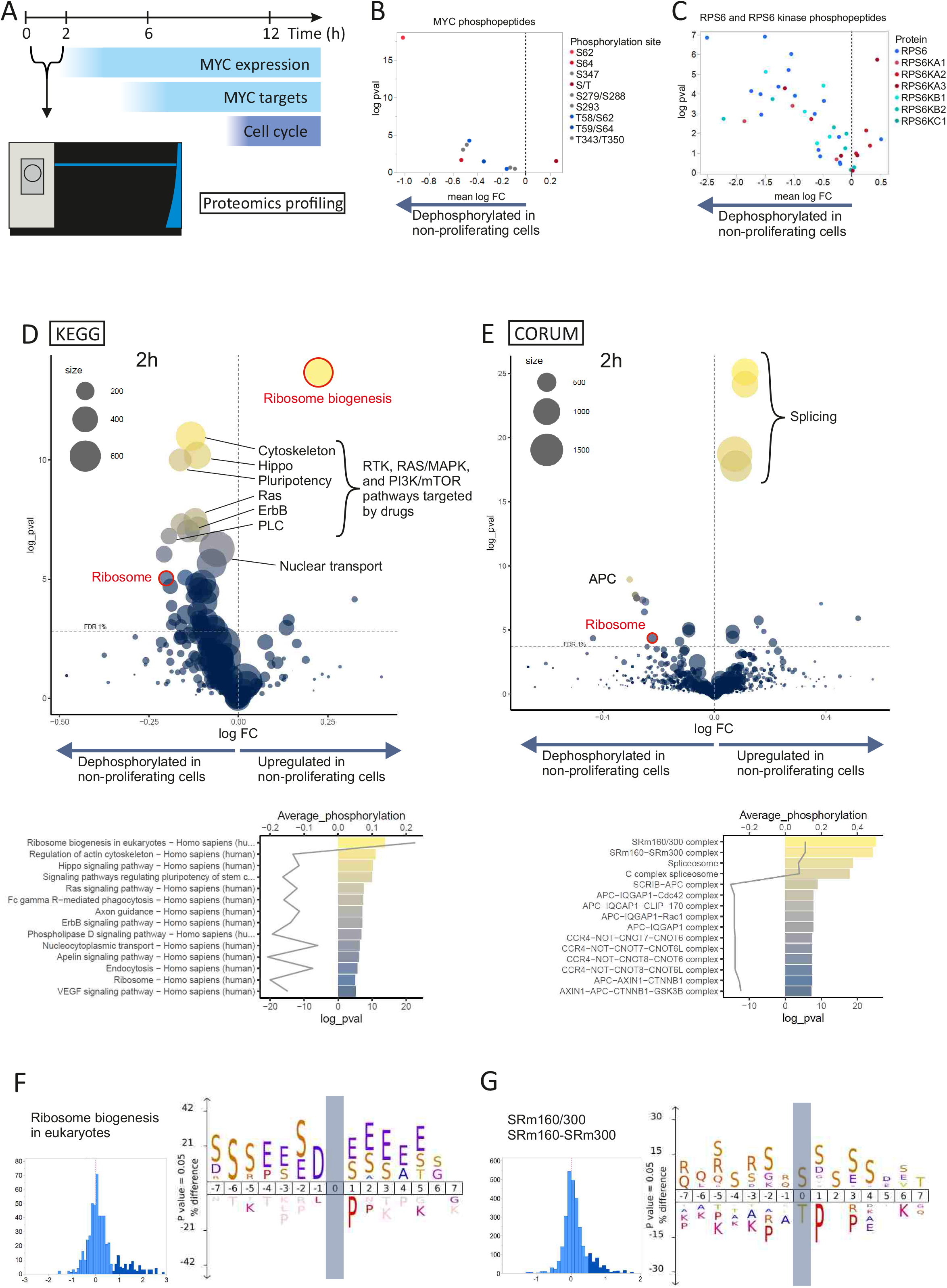
Phosphoproteomics profiling of shared targets of growth regulatory pathways. **A)** Phosphoproteomics analyses were performed in 30min and 2h timepoints before widespread transcriptional changes or cell cycle arrest **B)** Downregulation of MYC phosphopeptides in non-proliferating cells. In panels B-G, the data were generated from NCI-H1975, RKO, HCT116, LoVo (both resistance mechanisms), MCF-7 (Temsirolimus, both resistance mechanisms), and KBM7/HAP1 (both drugs) cell lines as presented in Fig. 2B. **C)** Downregulation of RPS6 and p70 S6K phosphopeptides in non-proliferating cells. **D)** Volcano plot: Average differential phosphorylation between proliferating and non-proliferating cells in all KEGG pathways excluding the KEGG DISEASE category is shown as log fold change on X-axis. p-value for one sample t-test is shown on Y-axis. Dashed line indicates 1% Benjamini-Hochberg FDR. Bar graph: Top pathways by p-value are indicated in the bar graph with colors corresponding to the volcano plot. Fold changes are shown with a line. **E)** Average differential phosphorylation between proliferating and non-proliferating cells in all CORUM database protein complexes is shown as log fold change on X-axis. p-value for one sample t-test is shown on Y-axis. Dashed line indicates 1% Benjamini-Hochberg FDR. Bar graph: Top complexes by p-value are indicated in the bar graph with colors corresponding to the volcano plot. Fold changes are shown with a line. **F)** Icelogo motif analysis of the upregulated phosphorylations (marked with darker blue in the histogram) compared to all phosphorylations (lighter blue) in the KEGG pathway ribosome biogenesis **G)** Icelogo motif analysis of the upregulated phosphorylations (marked with darker blue in the histogram) compared to all phosphorylations (lighter blue) in the CORUM database entries SRm160/300 and SRm160-SRm300

Analysis of common peptides whose phosphorylation was altered by drug only in the parental cell lines but not in the corresponding resistant derivative revealed multiple phosphorylation sites in proteins involved in metabolism, ribosome biogenesis and translation. Several known phosphoregulatory sites were identified, validating our approach. These included the well established phosphorylation of MYC and the ribosomal protein S6 and its upstream kinases (**Fig. 3B** and **C**). In addition, we found many shared downstream phosphorylation events, many of which affected essential genes or genes linked to cell proliferation (**Table S3**).

### Phosphorylation events target ribosome biogenesis prior to downregulation of gene expression

Due to the semi-random peptide detection inherent to mass spectrometry using data-dependent acquisition, individual phosphorylated peptides were commonly identified only in a subset of all the samples, potentially limiting the sensitivity of the analysis. To improve the sensitivity, we next identified increases and decreases of phosphorylation affecting entire protein complexes and pathways (see **Methods**). We observed increased phosphorylation in the large nucleolar proteins involved in rRNA processing and splicing (**Fig. 3D, E**). This upregulation occurred mostly in the acidophilic phosphorylation sites and serine clusters (**Fig. 3F, G**). In the case of ribosome biogenesis, these phosphorylations, occurring mainly in the Treacle protein (TCOF1), were short lived (**Fig. S3**). Ribosomal protein phosphorylation was reduced on average, and this change was also prominent at a later time point of 24 hours together with downregulated phosphorylation of ribosome biogenesis factors (**Fig S3**).

### Thermal proteome profiling identifies common targets in ribosome biogenesis and metabolism

We next performed thermal proteome profiling in three cell lines using proteome integral solubility-alteration assay (PISA;^23^). Consistently with the phosphorylation analysis, we detected a shift in thermal stability of ribosome biogenesis regulators after 2 h of drug treatment (**Fig. 4A, B**). We also detected effects on metabolic pathways (**Fig. 4A**). To further characterize the metabolic state of the cells after drug treatment, we performed metabolomics analysis using mass spectrometry. A targeted characterization of polar metabolites in selected cell lines 24h after drug treatment revealed a decrease in the nucleotide monophosphates UMP, AMP and GMP, and a block of the glycolytic pathway, based on accumulation of glucose and bisphosphoglycerate and decrease of fructose-6-phosphate (**Fig. 4C**). We also observed inhibition of *de novo* nucleotide synthesis, based on a decrease of levels of UMP, the key intermediate for synthesis of pyrimidine nucleotides via the *de novo* pathway (**Fig. 4C**, bottom right).

**Figure 4.**
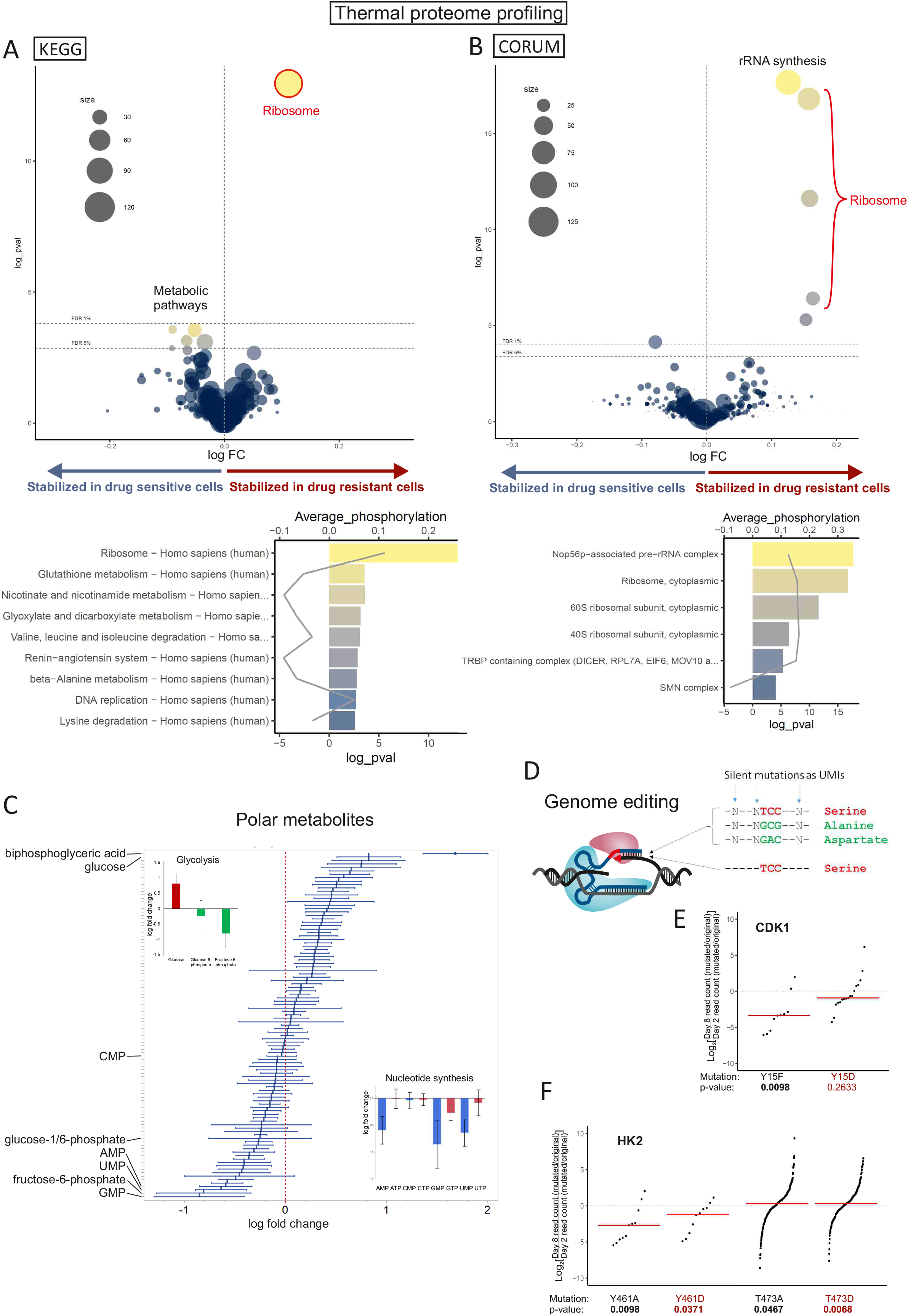
Protein interaction and metabolic changes in proliferating cells. **A)** Volcano plot: Average differential thermal stability change in PISA assay between drug resistant and sensitive cells in all KEGG pathways excluding the KEGG DISEASE category is shown as log fold change on X-axis. p-value for one sample t-test is shown on Y-axis. Dashed lines indicate 1% and 5% Benjamini-Hochberg FDR. Bar graph: Top pathways by p-value are indicated in the bar graph with colors corresponding to the volcano plot. Fold changes is shown with a line. In panels A and B, the data were generated from NCI-H1975, RKO, and HCT116 cell lines as presented in Fig. 2B, in 5 replicates for each condition. For each cell line pair, thermal stability change was calculated using the median of these replicates, and the statistic were calculated using the mean of these values. **A)** Volcano plot: Average differential thermal stability change in PISA assay between drug resistant and sensitive cells in all CORUM database protein complexes is shown as log fold change on X-axis. p-value for one sample t-test is shown on Y-axis. Dashed lines indicate 1% and 5% Benjamini-Hochberg FDR. Bar graph: Top pathways by p-value are indicated in the bar graph with colors corresponding to the volcano plot. Fold changes is shown with a line. **C)** Quantification of polar metabolites by LC-MS/MS. Mean +/- SEM for 8 cell line pairs is shown. Top left: Early glycolytic pathway metabolites glucose, fructose-6-phosphate, and fructose-1,6-bisphosphate. Bottom right: Nucleoside mono- and triphosphates. The data were generated from A549, NCI-H1975, RKO, HCT116, K562, MCF-7 (Palbociclib), T47D, and KBM7/HAP1 (both drugs) cell lines as presented in Fig. 2B **D)** CGE-assay. Phosphorylation sites were edited using prime editor or Cas9 with single stranded oligodeoxynucleotides (ssODN) as repair template for homology directed repair. For each editing reaction, pegRNA or ssODN library consisted of sequences that restore the phosphorylatable residue, introduce non-phosphorylatable residue (alanine or phenylalanine), and introduce phosphomimetic residue (aspartate or glutamate). Additional silent mutations were introduced to adjacent codons to enable lineage tracing. **E)** Consistently with previous publication^24^, the effect of mutating Y15 phosphorylation site in the CDK1 gene on fitness of HAP1 cells. For panels E and F, log2 values for day 8/day 2 ratios are shown for each sequence tag pair after calculating the ratio of read counts for mutated vs. original sequence at both timepoints. Red lines represent the median values, and p-values from Wilcoxon signed rank test are shown for each experiment. **F)** The effect of mutating Y461 and T473 phosphorylation sites in the HK2 gene on the fitness of HAP1 cells.

The accumulation of glucose suggested that the block in glycolysis could be due to decrease in activity of hexokinase, an enzyme that controls one of the rate-limiting steps of glycolysis. The gene for hexokinase2 (HK2) was also moderately downregulated by the chemotherapeutic drugs (**Fig. S4**). However, the relatively rapid changes in glycolytic metabolites, and decrease in thermal stability of components of metabolic pathways already at 2 hours (**Fig. 4A**) suggested that some effects on metabolism would be direct. We therefore tested whether phosphorylation of HK2 could explain the observed effects. We measured the requirement of known HK2 phosphorylation sites using competitive precision genome editing (CGE) assay (Fig 4D, E)^24^. We did not detect an effect on cell proliferation by mutation of HK2 residue T473, whose phosphorylation has been reported to affect HK2 activity ^25, 26^. However, this phosphorylation is not found in the publicly available mass spectrometry data sets, whereas phosphorylation in the nearby tyrosine residue (Y461) is commonly observed^27^. Mutation of Y461 in Hexokinase2 (HK2) inhibited growth of HAP1 cells (Fig. 4F), implicating phosphorylation of Y461 in regulation of HK2 activity. These results establish that transcriptional control and posttranslational regulation act in concert to regulate both ribosome biogenesis and metabolism.

### NOLC1 and TCOF1 define the proliferative compartment in human cancer

Our results described above establish that in cultured cells, the main process that associates with cell proliferation is ribosome biogenesis and translation. To determine whether this correlation is also observed in human tumors, we performed proteomic analyses of squamous cell carcinoma (SCC) of the tongue, a model that allows clear separation of proliferative and non-proliferative compartments of genetically similar tumor cells (**Fig. 5A**). In oral SCCs, cells in the inferior border of the tumor, the invasive front, have high proliferative activity ^28, 29^. Comparison of the invasive front and the central tumor enabled the identification of processes that are specific to proliferating tumor cells.

**Figure 5.**
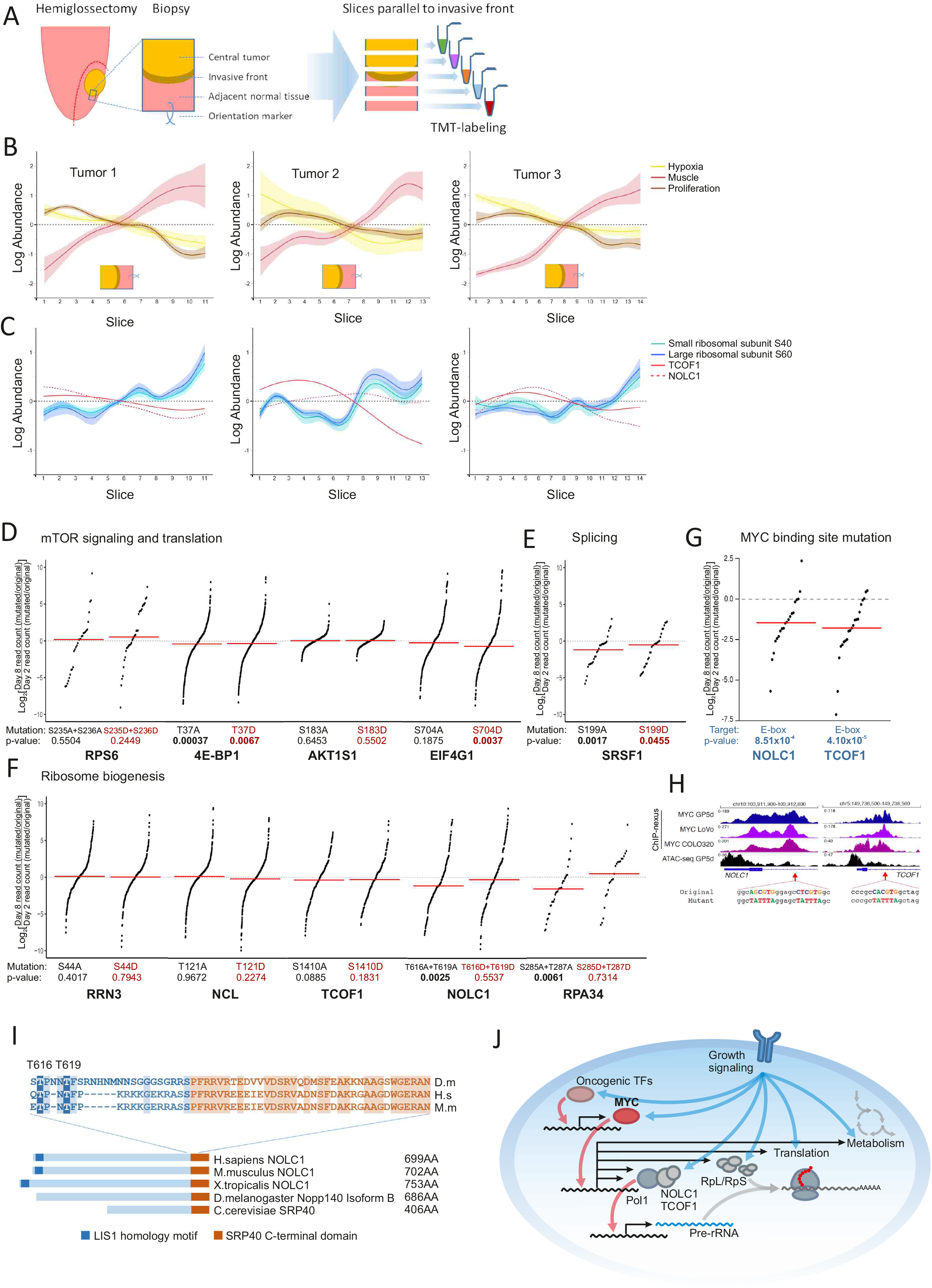
Fitness effect of selected phosphorylation site mutations and proliferation associated changes in head and neck squamous cell carcinoma (HNSCC) invasive front. **A)** HNSCC invasive front biopsies. Biopsies were collected from lateral tumors of the tongue following hemiglossectomy. Each biopsy was cut into 500µm slices parallel to the invasive front, and slices of the same tumor were analysed as one TMT-multiplex. **B)** Expression of hypoxia markers (GLUT1, PDK1), skeletal muscle specific proteins^52^, and proliferation markers (KI67, PCNA, MCM2-7) in the invasive front biopsies. Smoothing spline fit with 95% confidence interval shaded is shown. Tumor-normal tissue interface is marked by transition from proliferation and hypoxia marker expression to skeletal muscle specific protein expression. Proliferation marker expression is highest in the invasive front, whereas hypoxia marker expression is highest in the more central tumor. **C)** Expression of small and large subunit ribosomal proteins and ribosome biogenesis factors NOLC1 and TCOF1 in the invasive front. Smoothing spline fit is shown. For ribosomal proteins, 95% confidence interval for the fit is shaded. **D)** CGE-assay results for phosphorylation sites in effectors of mTOR signalling and translational regulators. In panels D-F, Log2 values for day 8/day 2 ratios are shown for each sequence tag pair after calculating the ratio of read counts for mutated vs. restored wild type sequence at both timepoints. Red lines represent the median values, and p-values from Wilcoxon signed rank test are shown for each experiment. **E)** The effect of mutating splicing factor SRSF1 phosphorylation site S199 on the fitness of HAP1 cells. **F)** The effect of mutating phosphorylation sites in ribosome biogenesis factors on the fitness of HAP1 cells **G)** The effect of mutating E-boxes in the promoters of NOLC1 and TCOF1 on the fitness of HAP1 cells **H)** Promoter regions of NOLC1 and TCOF1. MYC ChIP-nexus traces in three colorectal cancer cell lines, GP5d, LoVo, and COLO320 are shown in purple and ATAC-seq in GP5d cells in black. Targeted E-boxes and mutant sequences introduced by CGE are shown below the traces. **I)** Sequence and Interpro domain annotations of NOLC1 paralogs in H.sapiens, M.musculus. X.tropicalis, D.melanogaster, and S.cerevisiae. Alignment of the sequence consisting the identified proliferation associated phosphorylation sites T616/T619 and N-terminal half of the highly conserved SRP40 C-terminal domain in H.sapiens, M.musculus, and D.melanogaster. Conserved residues between fly and mammals are indicated with shaded background. Phosphorylations in the residues corresponding to human T616 and T619 have been observed in D.melanogaster and D.yakuba, respectively^32^ **J)** Schematic presentation of common targets of transcriptional and posttranslational growth regulatory mechanisms.

Biopsies were collected from tongue cancer patients treated with hemiglossectomy, which results in large surgical margin enabling the collection of a biopsy spanning the invasive front (**Fig. 5A**). Fresh frozen biopsies were cut into 500 µm sections parallel to the invasive fronts, enriching the invasive front cells to specific sections. Sections from one biopsy were analyzed as one TMT-multiplex, and protein expression specific to proliferating cells was identified by correlation profiling ^30^, where distributions of proteins across the slices were compared to known proliferation markers (KI-67, MCM2-7, PCNA) ^28, 29^. On the other hand, adjacent normal tissue exhibits high expression of proteins specific to skeletal muscle cells (**Fig. 5B**). TCOF1 and to a lesser extent NOLC1 exhibited a strong correlation with proliferation markers, whereas the ribosomal protein expression was highest in the adjacent normal tissue (**Fig. 5C**). This suggests that while both proliferating tumor cells and muscle cells require high protein synthesis capacity, the regulation of ribosome production has distinct characteristics in proliferating cancer cells, exemplified by elevated expression of TCOF1 and NOLC1.

### Phosphorylation of NOLC1 is required for cell proliferation

Our results establish that ribosome biogenesis correlates with cell proliferation both in cultured cells and in human tumors. The coordinated changes in phosphorylation and protein stability that correlate with cell proliferation suggest that at least some of the events might be mechanistically important for driving cell proliferation in response to phosphorylation signalling. To test this, we selected a set of phosphorylation sites for detailed analysis using CGE assay based on the following criteria: 1) essentiality of the protein, 2) conservation of the phosphorylation site, and 3) robust detection of the phosphopeptide in multiple different tumor types. We next measured the requirement of the phosphorylation sites using the CGE assay.

Control experiments established that some phosphorylation events that correlate with cell proliferation are not causative. For example, mutation of well-studied phosphorylation sites (S235 and S236) of ribosomal protein S6 had no effect, consistent with earlier observations ^31^ (**Fig. 5D**). Analysis of phosphorylation sites in other proteins involved in translation and mTOR signalling revealed also that a mutation of S183 in AKT1S1 had no effect. A minor effect was detected by mutation of another classical phosphorylation site, 4E-BP1 T37 (**Fig. 5D**). Interestingly, we found that a phosphomimetic mutation of S704 in EIF4G1 decreased cell proliferation (**Fig. 5D**), consistently with the upregulation of S704 phosphorylation in response to cytostatic drugs (**Fig. S5**).

Stronger effects were detected by mutating phosphorylation sites in proteins involved in two other processes, splicing (**Fig. 5E**) and ribosome biogenesis (**Fig. 5F**), that were rapidly modulated by cytostatic drugs. In particular, phosphorylation sites in two regulatory components of RNA polymerase I, NOLC1 and RPA34 had a strong negative impact on cell proliferation (**Fig. 5F**). Notably, both NOLC1 and its paralog TCOF1 were also regulated by MYC, and mutation of the MYC binding site in the promoter of either gene also caused a decrease in cell proliferation (**Fig. 5G, H**), establishing that NOLC1 regulation at both transcriptional and posttranslational level is required for cell proliferation. Notably, the phosphorylation of T616 in NOLC1 is conserved all the way to invertebrates such as *Drosophila*^*32*^ (**Fig. 5I**), suggesting that the identified mechanism of regulation is conserved across species.

## DISCUSSION

We find here that diverse types of upstream oncogenes, including the protein kinases and transcription factors that contribute to major forms of human cancer, converge to activate a single common downstream growth regulatory process. Our findings indicate that the gene regulatory network of cancer has an hourglass shape, with a large number of potential driver genes converging on a small number of master regulators, then diverging again into a large number of effector genes that underpin the cancer phenotype. Unlike the upstream drivers, which are typically tumor-type specific, the master regulators and their effector genes appear to be common to most, if not all, major forms of cancer.

The main processes activated by all the oncogenic mechanisms analyzed here are metabolism, translation, and ribosome biogenesis (**Fig. 5J**). Translation is affected both through phosphorylation of multiple components of the translation machinery, and by transcriptional control of multiple initiation factors by the master regulator MYC. Similarly, in ribosome biogenesis, the direct effects of phosphorylation signalling appear to affect proteins such as RPA34 and NOLC1 that are involved in ribosomal RNA synthesis, whereas the indirect action via MYC leads to upregulation of multiple proteins involved in ribosome biogenesis. Thus, the transcriptional control by MYC and posttranslational regulation by oncogenic kinase signalling act in concert to increase both the specific activity and concentration of components of protein synthetic machinery. In particular, we identify a key downstream node, NOLC1, whose activity is regulated both transcriptionally and posttranslationally. Reverse genetic experiments using CGE established that both types of regulation of NOLC1 are independently required for cell proliferation, highlighting the importance of this process (**Fig. 5J**).

The primary role of ribosome biogenesis for human cell proliferation and cancer is similar to simpler organisms such as *E coli* and yeast, where cell growth rate and ribosome concentration are linearly correlated ^33, 34^. Growth signals lead to an initial increase in ribosome biogenesis, followed by broader protein synthetic and anabolic activity. Our results suggest that multicellular organisms have evolved new control mechanisms to the old circuit that drives growth in response to nutrients. These mechanisms limit growth by hierarchical organization of cells into stem cells and differentiated cells ^35^, and by cell-to-cell signalling mechanisms that are required to specify or reinforce the proliferative state of specific cell types within particular tissues.

Initially, cancer was seen as a single disease, but more recent molecular analyses have revealed an extremely complex genetic origin for cancer. This has even led some investigators to conclude that cancer comprises many diseases that do not share a similarity of molecular mechanism. This view has lead to the development of “mechanism-based” antineoplastic agents that target specific upstream processes activated in particular tumor types, by for example directly binding to a single oncogenic protein kinase ^36^. Such targeted drugs are safe and effective, but in almost all cases lead to development of resistance due to mutations that prevent drug binding, or activate another upstream pathway driving the regrowth of the tumor. By contrast, our findings show that behind all the genetic complexity, there is a molecular commonality of mechanism shared by all major forms of human cancer.

This finding has very important implications to cancer therapy and prevention, as it suggests that it would be possible to develop a novel class of broad-spectrum antineoplastic agents that would not have the severe limitations of current chemotherapies. Existing antimetabolites and conventional chemotherapeutics are severely toxic, in part because they do not target specific proteins or pathways activated in tumors. Instead, they non-specifically damage DNA, and/or target multiple processes that are related to, but not identical to the mechanism uncovered here. For example, the commonly used chemotherapeutic 5-fluorouracil targets thymidylate synthase, an enzyme that acts in *de novo* thymidylate synthesis pathway, but it also causes DNA damage due to misincorporation of FdUTP into DNA^37^. Our results suggest that compounds that decrease the level of ribosome biogenesis or increase ribophagy, such as RNA polymerase I inhibitors^38, 39^, would more directly target a key consequence of oncogenic mutations. Furthermore, given the fact that genetic variants that decrease the activity of the pathway identified here decrease cancer risk but have no known harmful effect (**Fig. 1F**), a low dose or partial antagonist targeting the pathway might act as a chemopreventive agent, decreasing the incidence of a broad spectrum of human cancers.

## METHODS

### Cell lines

HAP1 (#C631) and KMB-7 (#C628) cell lines were obtained from Horizon Discovery and maintained in low-density cultures in Iscove’s Modified Dulbecco’s Medium (IMDM) according to the vendor’s guidelines. RKO, HCT116, LoVo, BT-474, T47D, MCF7, NCI-H1975, A549, CRL-2061, VCaP, and SK-N-MC cells were purchased from ATCC and cultured according to the vendor’s guidelines in the media specified by the vendor. To sensitize MCF-7 cells to temsirolimus, cells were cultured in reduced fetal bovine serum (5%) and without insulin. Prior to ChIP, MCF-7 cells were hormone starved for 48 h and subsequently mock-treated (minus ligand) or stimulated for 1 h with 100 nM Estradiol (E2). GP5d cells were obtained from Sigma (Sigma, 95090715), and cultured in DMEM supplemented with 10% fetal bovine serum, 2 nM L-glutamine and 1% penicillin– streptomycin. Wild-type and Myc-null Rat1 fibroblasts (Mateyak et al, 1997) were a kind gift from Professor John Sedivy, Brown University, and Professor Rene Bernards, Netherlands Cancer Institute. The cells were maintained in DMEM with 10% FBS and antibiotics.

PTEN knockout (RKO, HCT116, LoVo, BT-474, NCI-H1975, A549), RB1 knockout (T47D, MCF-7), NF1 knockout (MCF-7), and PIK3CA H1047R mutant (LoVo) cell lines were generated using Alt-R CRISPR-Cas9 from Integrated DNA Technologies according to vendor’s guidelines. Briefly, crRNA and tracrRNA duplex complexed with Cas9-HiFi protein was transfected using CRISPRMAX (Invitrogen). Transfection was verified after two days by imaging the fluorescence from the atto550 label in the tracrRNA. Single-stranded oligodeoxynucleotide (ssODN) was used as template homology-directed repair for PIK3CA H1047R mutation (Table S) at 3nM concentration. All crRNA sequences are listed in the Table S4. Resistant cells were selected by culturing with a drug concentration titrated to induce cell cycle arrest in the parental cell line. Single-cell colonies were cultured from cells that grew in the presence of the drug, and resistance mutation was verified by Sanger sequencing.

### ChIP-seq

Antibodies to Tcf4 (Clone 6H5-3 Exalpha Biologicals), β-catenin (Rabbit Polyclonal Antibody: H-102, Santa-Cruz Biotechnology), PAX3 (Rabbit Polyclonal Antibody Cat. No. CA1010, Calbiochem), GLI1 (Rabbit Polyclonal Antibody: H-300, Santa-Cruz Biotechnology), estrogen receptor (Rabbit Polyclonal Antibody: HC-20, Santa-Cruz Biotechnology), p300 (Rabbit Polyclonal Antibody: N-15, Santa-Cruz Biotechnology), RNA Pol II (Rabbit Polyclonal Antibody: H-224, Santa-Cruz Biotechnology), α-H3K4me1 (Rabbit Polyclonal Antibody: ab8895, Abcam) and normal IgG (mouse: sc-2025; rabbit: sc-2027, Santa-Cruz Biotechnology). were used in ChIP. Chromatin immunoprecipitation by sequencing was performed as described in Tuupanen et al.^40^

The ChIP-seq data was analyzed as described in Wei et al.^41^ Sequencing reads were mapped to the NCBI36 release of the human genome using Maq version 0.6.5.^42^ Only reads of mapping quality score ≥ 30 were accepted. Reads were then extended to estimated fragment length, and peak height was determined at each position as the number of overlapping extended reads. For each peak of height eight or more, the total number of sequences in the continuous region of four or more overlapping sequences was compared to the number of sequences in the same region in the IgG control. The probability of observing the difference between the sequence counts in the ChIP sample and IgG control by chance was estimated using the Winflat program^43^ and peaks that had probability smaller than 0.05 (without multiple hypothesis testing correction) were selected for further analysis.

### siRNA treatment and expression profiling

For siRNA knockdown, predesigned FlexiTube siRNAs (QIAGEN) for the TFs targeted in ChIP, as well as control siRNA (SI03650325, QIAGEN), were transfected into cells using HiPerFect Transfection Reagent (Cat. 301704, QIAGEN). Transfections were performed in two steps with 24h interval, and the cells were harvested 48h after the transfections. Total RNA were prepared by QIAshedder (Cat. 79656, QIAGEN) and Qiagen RNAeasy Kit (catalog no. 74104). Expression profiling was performed using Affymetrix human genome U133plus2.0 arrays. The background noise was then corrected according to the GCRMA method^44^. The probesets were subsequently filtered for expression measurements to have a value of 100 fluorescence units in at least 25% of the samples.

Differential expression analysis was performed by fitting a linear model as described in the Limma package^45^; p values were adjusted according to Benjamini and Hochberg’s method to control the false discovery rate. Probes showing significant differences (p < 0.01) between control samples were eliminated from the analysis.

### GWAS variant information

All SNPs that were associated either with the trait “cancer” or a trait that had “cancer” as a parent trait were downloaded from the GWAS catalog (date 2021-11-19; ^46^). There were 7,796 SNPs associated with at least one of the cancer traits and 11,155 cancer trait associations with one of the SNPs.

### Single-cell RNA-sequencing

CD298 and β2M antibodies from Biolegend () were labeled with TotalSeqB-oligos (Table S) according to a protocol by Van Buggenum et al.^47^ Cells were cultured on 24 well plates and treated with varying drug concentrations and durations. In the end of the drug treatment, each row and column on the plate was labeled according to the cell hashing protocol by Satija lab (https://cite-seq.com/protocols/) with a unique TotalSeqB barcode resulting in unique combination of two barcodes for each treatment condition. After the labeling, cells from the whole plate were pooled and resuspended in PBS, and sequencing libraries were prepared on a Chromium platform using Single Cell 3’ v3 and feature barcoding reagent kits (10x Genomics, Pleasanton, CA) according to manufacturer’s instructions. For each plate, 2500-5000 cells were sequenced at a depth of 50000-100000 reads per cell on a NovaSeq (Illumina). Preprocessing of the data was performed using CellRanger (10x Genomics). Cells were further filtered based on read count and TotalSeqB barcode count distribution that could not be clearly assigned to a treatment condition to remove probable empty beads. Gene expression was modeled as a function of drug concentration, treatment duration, cell cycle phase and presence of resistance mutation, and genes were ranked by the explanatory power of the drug effect in the model.

### Phosphoproteomics

Cell line samples were collected at early (30min or 2h) and late (24h) treatment timepoints. The cells were washed twice with ice cold PBS and scraped into cold PBS with phosphatase inhibitors (PhosSTOP, Roche). Pelleted cells were snap-frozen and lysed with 8M urea buffer with 100nM TEAB (Sigma). Lysates were reduced with DTT at final concentration of 20mM and alkylated with IAA at final concentration of 40mM. For digestion, the buffer was diluted to <2M urea concentration, and samples were digested over night with Lys-C (Wako) followed by 4h digestion with Trypsin (Promega) at room temperature. Digests were desalted and 200-400µg of each sample was labeled with TMT or TMTPRO reagents. Multiplexed samples were reverse-phase fractionated at high pH on an Acquity UPLC system (Waters) similar toChristoforou et al.^48^ 10% of each fraction was taken for total protein analysis and the remaining was enriched with a modified SIMAC procedure. Briefly, dried peptides were reconstituted 50% ACN, 0.1% TFA, and incubated with TitanSpehere beads (Hichrom Ltd) loaded in equal volume in 80% Acetonitrile, 5% TFA and 1 M Glycolic acid for 30 minutes with shaking (≥4:1 beads:peptides ratio). Beads were washed once with 80% ACN, 1% TFA, once with 10% ACN, 0.1% TFA, and eluted with ∼1.2% ammonia after 10 minute incubation with shaking. Supernatants from loading and wash steps were pooled for 5-6 fractions, and each pool was dried and enriched with High-Select™ Fe-NTA Phosphopeptide Enrichment Kit (ThermoFisher Scientific).

The LC-ESI-MS/MS analyses were performed on a nanoflow HPLC system (Easy-nLC1000, Thermo Fisher Scientific) coupled to the Orbitrap Fusion Lumos mass spectrometer (Thermo Fisher Scientific, Bremen, Germany) using synchronous precursor selection SPS-MS^3^ acquisition method. Mass spectrometry methods are described in detail in Supplementary Methods

Frozen tumor biopsies were cut into 500µm slices, and each slice was pulverized with a dounce in a microcentrifuge tube. Pulverized samples were lysed, reduced, alkylated and desalted similar to cell line samples. 100µg of each slice was labeled with TMTPRO, and pooled sample was off-line-fractionated with Agilent 1260 HPLC system following a protocol by Mertins et al.^49^ Fractions were pooled to 12 samples and 10% of each fraction was taken for total protein analysis. Remaining sample was enriched with High-Select™ Fe-NTA Phosphopeptide Enrichment Kit (ThermoFisher Scientific). The LC-ESI-MS/MS analyses were performed on a nanoflow HPLC system (Easy-nLC1200, Thermo Fisher Scientific) coupled to the Orbitrap Fusion Lumos mass spectrometer (Thermo Fisher Scientific, Bremen, Germany) equipped with a nano-electrospray ionization source and FAIMS interface (Thermo Fisher Scientific).

### Protein interaction analysis by PISA assay

PISA assay was performed as described previously^23^. Samples were collected at 2h and 24h treatment time points. Each resistant/sensitive cell line pair was analyzed in 5 replicates for drug treated and control cells split between two TMT 11-plexes with a calibrator sample. TMT-multiplexes were fractionated to 24 fractions as described previously^23^ and analyzed on Thermo Orbitap Q-Exactive HF coupled to UltiMate 3000 RSLC nanoUPLC system. The data were normalized by median, and the data between different TMT-multiplexes were combined by calculating a ratio to the calibrator sample included in both multiplexes. Drug induced changes were identified by comparing the treated cells to non-treated cells, and proliferation associated changes were identified by comparing the drug induced changes between sensitive and resistant cells.

### Metabolite analyses

Samples were collected after 24 hour drug treatment, and intra-cellular metabolites were extracted using a methanol/chloroform method. Briefly, 600 µl of methanol/chloroform (2:1 v/v) was added to each cell pellet (∼50 µl). Samples were vortex mixed and sonicated for 15 minutes. Two hundred microliters of chloroform and water were added, and samples were vortex mixed and separated by centrifugation at 17000g for 15 minutes. The aqueous and organic layers were then collected, and the procedure was repeated using halved volumes on the residual sample containing the precipitated protein to maximize the recovery. Collected layers from two round of extraction were combined, and the aqueous layer was dried overnight in vacuum centrifuge (Eppendorf), while the lipid fraction was dried under nitrogen gas flow. Dried samples were stored at -80 °C.

Targeted metabolite analysis was performed with reverse and normal phase separations on a Thermo scientific UHPLC^+^ series coupled with a TSQ Quantiva mass spectrometer in operated in positive and negative ion mode at the same time.

### Fitness effect of phosphorylation sites and MYC target sites by CGE assay

CGE assay was performed as described in Pihlajamaa et al^24^. Briefly, 200,000-400,000 early-passage HAP1 cells were transfected with a RNP complex containing tracrRNA and crRNA (250-500ng) with S.p. HiFi Cas9-protein (1-2µg; Integrated DNA Technologies) together with ssODN HDR template with a final concentration of 3 nM using CRISPRMAX (Life Technologies) following manufacturer’s recommendations. For prime-editor experiments, plasmids for prime-editor and pegRNA, pCMV-PE2 and pU6-pegRNA-GG-acceptor (Addgene #132775 and #132777, respectively)^50^, were transfected using FuGENE HD (Promega) according to manufacturer’s instructions and 4:1 Fugene HD:DNA ratio. Half of the cells were collected for gDNA isolation on day 2 after transfection, and the other half cultured until day 8 for late timepoint sample. Isolation of gDNA, treatment with RNase A, exonucleases I and VII, two-step PCR amplification, sequencing, and data analysis were performed as described in Pihlajamaa et al^24^. Read count 5-50 was used as a cutoff for day 2 samples depending on the sequencing depth. Sequences of crRNAs, HDR donor templates, pegRNAs, and target-specific primers are listed in the Table S4.

### Patient material

Collection of HNSCC invasive front samples was approved by the Ethics Committees of Southwest Finland (ETMK 166/1801/2015) and Turku University Central Hospital (TO6/022/17) and was conducted according to the principles of Declaration of Helsinki. Biopsies were taken from the resected tumors without compromising the routine diagnostics, and the orientation of the biopsy was marked with a stich prior to snap-freezing the sample.

